# Laser-facilitated epicutaneous immunotherapy with hypoallergenic beta-glucan neoglycoconjugates suppresses lung inflammation and avoids local side effects in a mouse model of allergic asthma

**DOI:** 10.1101/2020.01.18.911123

**Authors:** Evgeniia Korotchenko, Victoria Schießl, Sandra Scheiblhofer, Isabella Joubert, Helen Strandt, Theresa Neuper, Muamera Sarajlic, Renate Bauer, Mark Geppert, David Joedicke, Sabrina Wildner, Susanne Schaller, Stephan Winkler, Gabriele Gadermaier, Jutta Horejs-Hoeck, Richard Weiss

**Affiliations:** University of Salzburg, Department of Biosciences, Salzburg, Austria; University of Applied Biosciences Upper Austria, Research and Development, Hagenberg, Austria

## Abstract

**Background:** Allergen-specific immunotherapy via the skin targets an area rich in antigen presenting cells, but can be associated with local and systemic side effect. Allergen-polysaccharide neoglycogonjugates can increase immunization efficacy by targeting and activating dendritic cells via C-type lectin receptors and reduce side effects.

**Objective:** We investigated the immunogenicity, allergenicity and therapeutic efficacy of laminarin-ovalbumin neoglycoconjugates (LamOVA).

**Methods:** The biological activity of LamOVA was characterized *in vitro* using bone marrow derived dendritic cells. Immunogenicity and therapeutic efficacy was analyzed in BALB/c mice. Epicutaneous immunotherapy (EPIT) was performed using fractional infrared laser ablation to generate micropores in the skin and the effects of LamOVA on blocking IgG, IgE, cellular composition of BAL, lung, and spleen, lung function, and T cell polarization was assessed.

**Results:** Conjugation of laminarin to ovalbumin reduced its IgE binding capacity 5-fold and increased its immunogenitiy 3-fold in terms of IgG generation. EPIT with LamOVA induced significantly higher IgG levels than OVA, matching the levels induced by s.c. injection of OVA/alum (SCIT). EPIT was equally effective as SCIT in terms of blocking IgG induction and suppression of lung inflammation and airway hyperresponsiveness, but SCIT was associated with a higher level of therapy induced IgE and TH2 cytokines. EPIT with LamOVA induced significantly lower local skin reactions during therapy compared to unconjugated OVA.

**Conclusion:** Conjugation of the allergen to laminarin increased its immunogenicity while at the same time reducing local side effects. LamOVA EPIT via laser generated micropores is safe and equally effective to SCIT with alum, without the need for adjuvant.

## Introduction

Epicutaneous allergen-specific immunotherapy (EPIT) has been introduced almost 100 years ago and only recently revisited ^1^. Current studies show that EPIT can lead to a potent reduction of allergic symptoms and that the efficacy of the therapy is dependent on the allergen dose and the pretreatment of the skin. However, depending on the degree of barrier disruption, epicutaneous application of allergen can induce local and even systemic side effects in some individuals ^2^. Alternatively, therapy has been performed by application of occlusive patches to intact skin leading to epicutaneous absorption due to maceration ^3^. The major pitfall of this method is the inefficient uptake, requiring numerous applications of high doses ^4^.

Due to the high number of antigen presenting cells (APCs) in the epidermis and dermis, skin based immunization often requires smaller amounts of antigen to induce immune responses compared to subcutaneous (s.c.) or intramuscular injections ^5^. APCs express a large variety of pattern-recognition receptors (PRRs) determining their main function, i.e. sensing of pathogens and induction of immune reactions ^6^. One group of PRRs are C-type lectin receptors (CTLs), which bind to carbohydrates in a calcium-dependent manner. In cohort with other receptors, stimulation of CTLs determines the activation status of dendritic cells (DCs) and subsequent T cell polarization ^7^. Therefore, the use of protein neoglycoconjugates for immunizations can increase vaccine potency.

As a prominent member of the CTL family, Dectin-1 is functionally equivalent in mice and humans ^8^. It recognizes β-glucans and some proteins such a tropomyosin ^9^, and upon ligand binding activates innate immune responses. Murine dectin-1 is expressed on macrophages, neutrophils and dermal dendritic cells ^10^. In humans, dectin-1 is not restricted to cells of the myeloid lineage but can also be found on epithelial cells ^8, 11, 12^, keratinocytes ^13^, B cells, and subpopulations of T cells ^8^. As dectin-1 is expressed on dermal DCs, its use as a vaccination platform for targeted skin vaccination is highly promising. Here, we used 6kDa laminarin/ovalbumin (LamOVA) conjugates for allergen specific immunotherapy (AIT). Laminarin has strong immunostimulatory properties and has been employed in tumor therapy for activation of the innate immune system ^14^. Moreover, polysaccharides can mask epitopes on allergens preventing IgE cross-linking on mast cells and histamine release, decreasing the risk of side effects ^15^.

Skin has a protective function, and its outermost layer, the stratum corneum, represents a tight barrier that has to be overcome for antigen delivery. We use infrared laser to form micropores in the upper layers of the skin ^16^. Besides facilitating vaccine penetration, the local tissue damage caused by laser treatment attracts large numbers of APCs and creates a pro-inflammatory milieu, thereby acting as a physical adjuvant for skin vaccinations ^17^.

In our current work we evaluated the immunostimulatory capacity of LamOVA conjugates and their efficacy in a preventive and therapeutic BALB/c mouse model of allergic lung inflammation. Laser-facilitated EPIT with LamOVA significantly reduced airway hyperresponsiveness and local side effects *in vivo*.

## Materials and Methods

A more detailed version of material and methods can be found in the online supplement.

### Generation and analysis of allergen-carbohydrate neoglycoconjugates

6 kDa laminarin from laminaria digitata (Sigma, L9634) was dialyzed with a cut-off of 2 kDa to remove low molecular weight impurities, partially oxidized (20%) with sodium (meta)periodate and coupled to endotoxin-free ovalbumin (OVA, EndoFit™ Ovalbumin, InvivoGen) using 2-step reductive amination with 50 mM sodium cyanoborohydride (NaCNBH_3_) and 10 mM sodium borohydride (NaBH_4_) as previously described ^15^. Resulting neoglycoconjugates were separated by size-exclusion chromatography on a HiPrep 26/100 Sephacryl^®^ S-200 HR column (GE Healthcare Life Sciences), and fractions were analyzed by 10% SDS-PAGE. Fractions containing high molecular weight (>70kDa) laminarin-ovalbumin conjugates (LamOVA) were used for further experiments.

Carbohydrate concentration in conjugates was measured using anthrone method ^18^ and OVA concentration was determined by amino acid analysis ^19^. The hydrodynamic radius of conjugates was analyzed by dynamic light scattering using a Zetasizer Nano ZS with a DTS1070 capillary cell (Malvern Instruments).

Biological activity of laminarin in LamOVA conjugates was analyzed by ELISA using a soluble murine Fc-Dectin-1a receptor (InvivoGen). Hypoallergenicity was analyzed *in vitro* as a measure of rat basophil leukemia cell (RBL-2H3) activation by β-hexosaminidase release assay ^15^.

### Uptake and activation analysis using bone marrow-derived dendritic cells

Bone marrow-derived dendritic cells (BMDCs) were harvested from mouse femur and tibia and incubated with either 20ng/mL murine GM-CSF (Immunotools, Cat.Nr. 12343125) or 200ng/mL human Flt3-L (AcroBiosystems, Cat.Nr. FLL-H55H7) as described ^20, 21^ with minor changes.

LamOVA effects on CD86 and Dectin-1 expression on CD11c+ Flt3-L – generated DCs (FL-DCs) and on CD11c+ MHCII^high^ CD11b^int^ GM-CSF – derived DCs (GM-DCs) were assessed. Activation of OVA-specific naïve DO11.10 T cells was analyzed in co-culture assays. BMDC activation, T cell proliferation and uptake were assessed by flow cytometry as described in supplementary materials. Cytokine concentration in supernatants was analyzed by LEGENDplex assay (Biolegend) using the 13-plex mouse inflammation panel and the 13-plex mouse proinflammatory chemokine panel according to the manufacturer’s instructions.

### Animal experiments

Female BALB/c mice aged 6-8 weeks were obtained from Janvier (Le Genest-Saint-Isle, France) and maintained at the animal facility of the University of Salzburg in an SPF environment according to local guidelines. All animal experiments were conducted in compliance with EU Directive 2010/63/EU and have been approved by the Austrian Ministry of Education, Science and Research, permit number BMBWF-66.012/0006-V/3b/2018. Blood samples were drawn on days 27, 35, 55 (prevention) and 16 and 86 (therapy) by puncture of v. saphena.

To study immunogenicity of the conjugates, mice (n=6) were immunized three times with 20 µg of OVA, LamOVA conjugates (20 µg OVA coupled to 49µg laminarin), a mixture of equivalent amounts of OVA + laminarin, or laminarin and PBS (sham controls). The antigens were applied to laser-microporated skin on days 1, 15, and 29. Microporation was performed using a P.L.E.A.S.E. professional infrared laser device (Pantec Biosolutions, Ruggell, Liechtenstein) with a total fluence of 8.3 J/cm^2^ as previously described ^15^. After the third immunization, mice were sensitized two times in a ten-day interval (days 36 and 46) by i.p. injections of 1 µg EndoFit™ OVA adsorbed to 50% (v/v) alum (Alu-Gel-S, Serva) in endotoxin-free PBS. To induce lung inflammation, animals were challenged by exposure to aerosolized grade V OVA from Sigma (A5503), that has been purified from endotoxin using Triton-X114 method ^22^ (10 mg/mL in 0.9% NaCl, LPS < 0.39 pg per 1 µg OVA), for 30 min on days 53-55 using a Pariboy SX jet nebulizer with a Pari LL nebulizer head (Pari GmbH).

To study therapeutic efficacy, animals (n=12) were sensitized by two i.p. injections with 10 µg of EndoFit™ OVA/alum on days 1 and 8. On day 15, mice were challenged intranasally with 10 µg of EndoFit™ OVA in 40 µL PBS, under isoflurane anesthesia. One day after the challenge, lung function and specific serum IgE were analyzed to confirm that all animals in the treatment groups had an asthma-like phenotype and OVA-specific IgE. Eight therapeutic immunizations were performed between days 18 and 67 in weekly intervals. Mice were treated with LamOVA conjugates, OVA, laminarin, or PBS. 35µg of OVA and/or 86µg of laminarin equivalents were applied on 1 cm^2^ of laser-microporated skin. As a positive control, s.c. injections with 35 µg EndoFit™ OVA/alum were performed. On days 84-86, mice were challenged with OVA aerosol.

### Analysis of local side effects

For analysis of erythema severity, mice were photographed on days 47 or 54 with a SONY ILCE-7M2 camera, 28-70 mm zoom lens with a constant source of light using .JPG format. MATLAB R2019a was used for analysis. Briefly, the area of laser microporation (region of interest, ROI) was defined by an investigator blinded to the treatment. Based on the grey value of the selected ROI, a threshold was manually selected, to define the erythema that was darker than healthy skin. Erythema size and mean blob size were calculated from the selected ROI. Erythema size is calculated as the ratio of pixels above the chosen threshold to the number of pixels inside the ROI. A blob is defined as the sum of directly connected pixels of the segmented erythema. The mean value of the five biggest blobs was evaluated.

### Lung function analysis

The day after the aerosol challenge, lung function was analyzed by whole body plethysmography (WBP) using a Buxco 6-chamber unrestrained WBP system (Data Sciences International). The next day, lung resistance/compliance measurement was performed using a FinePointe Series Resistance and Compliance (RC) site (Data Sciences International). Bronchoalveolar lavage fluid (BALF) was collected and its cellular composition was analyzed by flow cytometry staining for Siglec F, CD45, CD4, CD8, CD19, and Ly6G/Ly6C (Gr-1).

### Lung digestion

Lung tissue was digested and total leukocytes were separated using biotinylated CD45 and BD™ IMag Streptavidin Particles (BD Biosciences). Cells were stained for CD11b, CD24, CD4, CD45, Ly6G/Ly6C (Gr-1), MHC-II, and Siglec F and analyzed by flow cytometry.

### Analysis of OVA specific serum IgG/IgE and cell-bound IgE

OVA-specific IgG isotypes in sera were analyzed by direct ELISA. OVA-specific serum IgE was measured using a rat basophil leukemia cell (RBL-2H3) assay. Briefly, cells were incubated for 2h with mouse sera diluted 1:150 in medium, followed by three washing steps and 1 h stimulation with 0.3µg/mL EndoFit™ OVA. β-hexosaminidase levels in supernatants were measured and degranulation was calculated as percentage of total lysis from control wells lysed with Triton X-100.

Cell-bound IgE was analyzed *ex vivo* by basophil activation test from whole blood drawn one day after the second aerosol challenge. Cells were stimulated in the presence or absence of autologous serum with 2 ng/mL OVA for 2h at 37°C, 5% CO_2_, 95% humidity. After the incubation, cells were stained for IgE, CD4, CD19 and CD200R and analyzed by flow cytometry on a FACS Canto II flow cytometer (BD Biosciences).

### Lymphocyte restimulation

Spleens were harvested for T-cell restimulation with OVA. After erythrocyte lysis in Ammonium-Chloride-Potassium (ACK) lysing buffer, splenocytes were resuspended in T-cell medium (RPMI-1640, 10% FCS, 25mM Hepes, 2 mM L-Glu, 100 µg/mL streptomycin, 100U/mL penicillin) and 0.6×10^6^ cells/well were stimulated with 0.1 mg/mL EndoFit™ OVA and incubated for 3 days at 37°C, 5% CO_2_. Supernatants were harvested and analyzed by LEGENDplex immunoassay using the mouse T helper 13-plex cytokine panel. Cells were stained for CD4, CD25, CD44, FoxP3, GATA3 and analyzed by flow cytometry on a Cytoflex S flow cytometer (Beckman Coulter).

### Statistical analysis

Data was statistically analyzed using GraphPad Prism 7 using one-way ANOVA with Tukey’s post-hoc test unless otherwise specified. P-values are shown as: * P≤ 0.05, ** P ≤ 0.01, *** P ≤ 0.001, **** P ≤ 0.0001.

## Results

### Generation of laminarin OVA conjugates

Neoglycoconjugates were generated by coupling of 6 kDa polysaccharide laminarin to EndoFit™ OVA via mild periodate oxidation and reductive amination. Size-exclusion chromatography (suppl. Fig. 1A) was used to separate neoglycoconjugates by size, followed by SDS-PAGE of the fractions (suppl. Fig. 1B). Based on visual analysis, fractions bigger than 70 kDa were pooled (LamOVA) and further analyzed.

To determine OVA concentration in conjugates, amino acid analysis was used (suppl. Fig 1C), as common methods of protein quantification such as UV absorption, Bradford assay, or BCA assay are all affected by the carbohydrate moiety. Additionally, this method allows estimating the amount of lysine residues which participated in reductive amination. For LamOVA, 37.5% of lysins were coupled to laminarin. Concentration of laminarin was assessed by anthrone method, and carbohydrate/protein ratio was 2.5.

Conjugates were homogeneous in size and had a diameter of 8.27±0.75nm as shown by dynamic light scattering (suppl. Fig. 1D).

### Laminarin is biologically active after conjugation

To ensure that the structure of laminarin was not destroyed during mild periodate oxidation and that the polysaccharide remained biologically active after coupling, binding to Dectin-1 was assessed by ELISA. As shown in Fig. 1A, LamOVA was recognized by Dectin-1, whereas no binding of unconjugated OVA was observed.

**Fig. 1.**
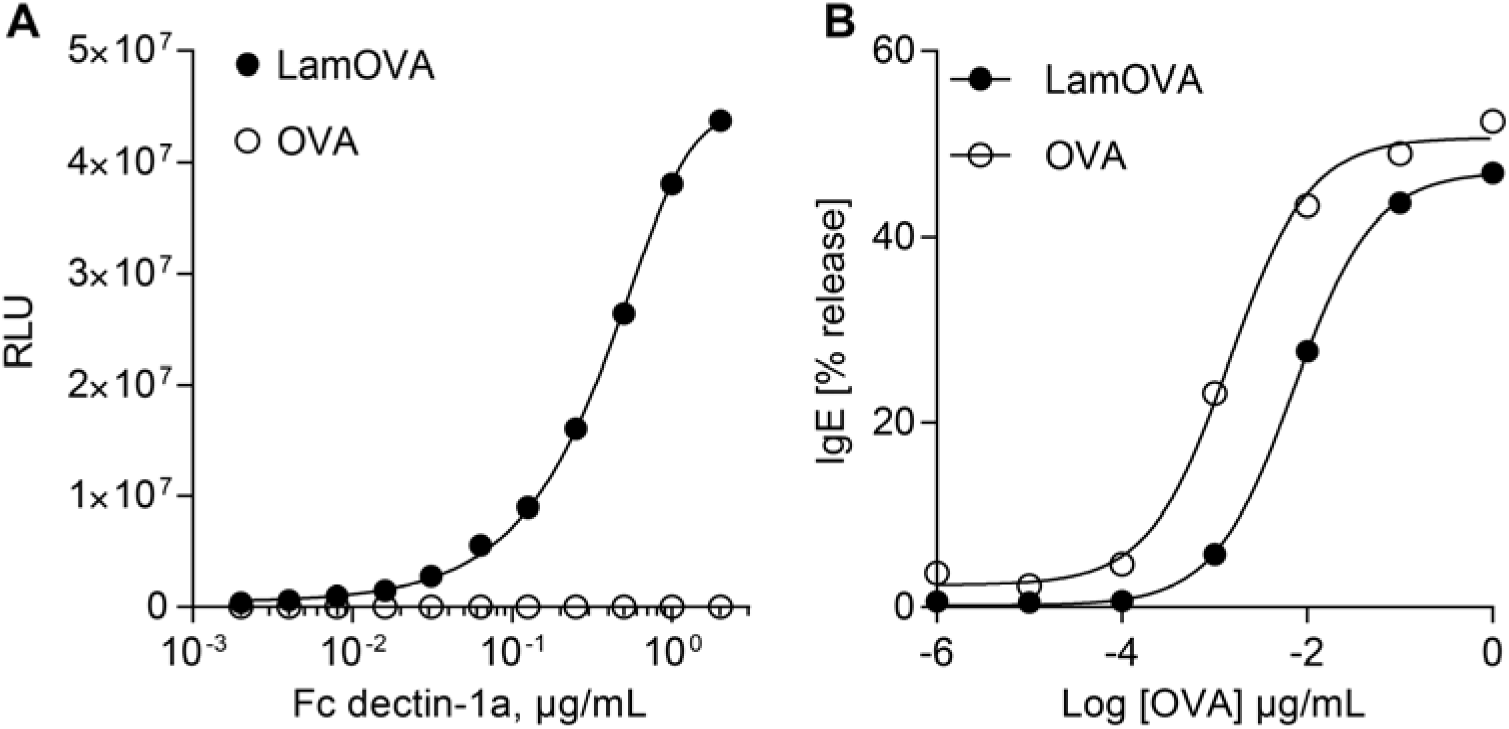
A) Binding of soluble dectin-1a receptor (fused to IgG-Fc) to plate-bound LamOVA or OVA. Data are shown as relative light units (RLU) of a luminometric ELISA. B) Stimulation of IgE-loaded RBL cells with LamOVA or OVA. Basophil IgE-mediated degranluation is presented as percentage of total beta-hexosaminidase release.

### LamOVA conjugates activate basophils to a 5-fold lesser extent than unmodified OVA

Hypoallergenicity is a prerequisite for AIT via the skin, as administration of native allergen to barrier disrupted skin has been shown to induce local or even systemic side effects ^2^. We hypothesized that laminarin polysaccharide chains (6 kDa) would mask IgE epitopes of OVA and prevent mast cell and basophil degranulation *in vivo*. To prove that conjugates were less allergenic than unconjugated OVA, rat basophil leukemia (RBL) cells were incubated with sera of highly sensitized mice and stimulated with LamOVA or OVA. Beta-hexosaminidase secretion after antigen stimulation is a measure of IgE cross-linking. As shown in Fig. 1B, a 5-fold higher dose of LamOVA was required to activate IgE loaded basophils compared to OVA thus confirming hypoallergenicity of LamOVA.

### Laminarin conjugation to OVA facilitates uptake by BMDCs and induces their activation

Bone marrow cells were incubated with either hFLT3-L (FL) or mGM-CSF (GM) to generate BMDCs. Uptake of pHrodo-labeled LamOVA was significantly enhanced compared to pHrodo-OVA in both FL-BMDCs and GM-BMDCs (Fig. 2A). LamOVA significantly activated BMDCs in a dose-dependent manner, whereas unconjugated OVA and laminarin (used at doses equivalent to those present in LamOVA) did not induce CD86 expression even at the highest doses. GM-BMDCs showed a higher base-line expression of CD86 and responded more strongly to positive control stimulus LPS than to LamOVA (Suppl. Fig. 2A). In contrast, FL-BMDCs had a more naïve phenotype and LamOVA induced an equivalent upregulation of CD86 compared to LPS (Suppl. Fig. 2B). In both cell types, LamOVA induced significantly higher CD86 expression than OVA, laminarin and PBS stimulation (Suppl. Fig. 2A and B). LamOVA and laminarin stimulation significantly downregulated dectin-1 surface expression in GM-BMDCs and FL-BMDCs (Suppl. Fig. 2C and D) indicating binding of the beta-glucan to its receptor. Interestingly, LPS activation also induced downregulation of dectin-1, which has already been observed in macrophages ^23^.

**Fig. 2.**
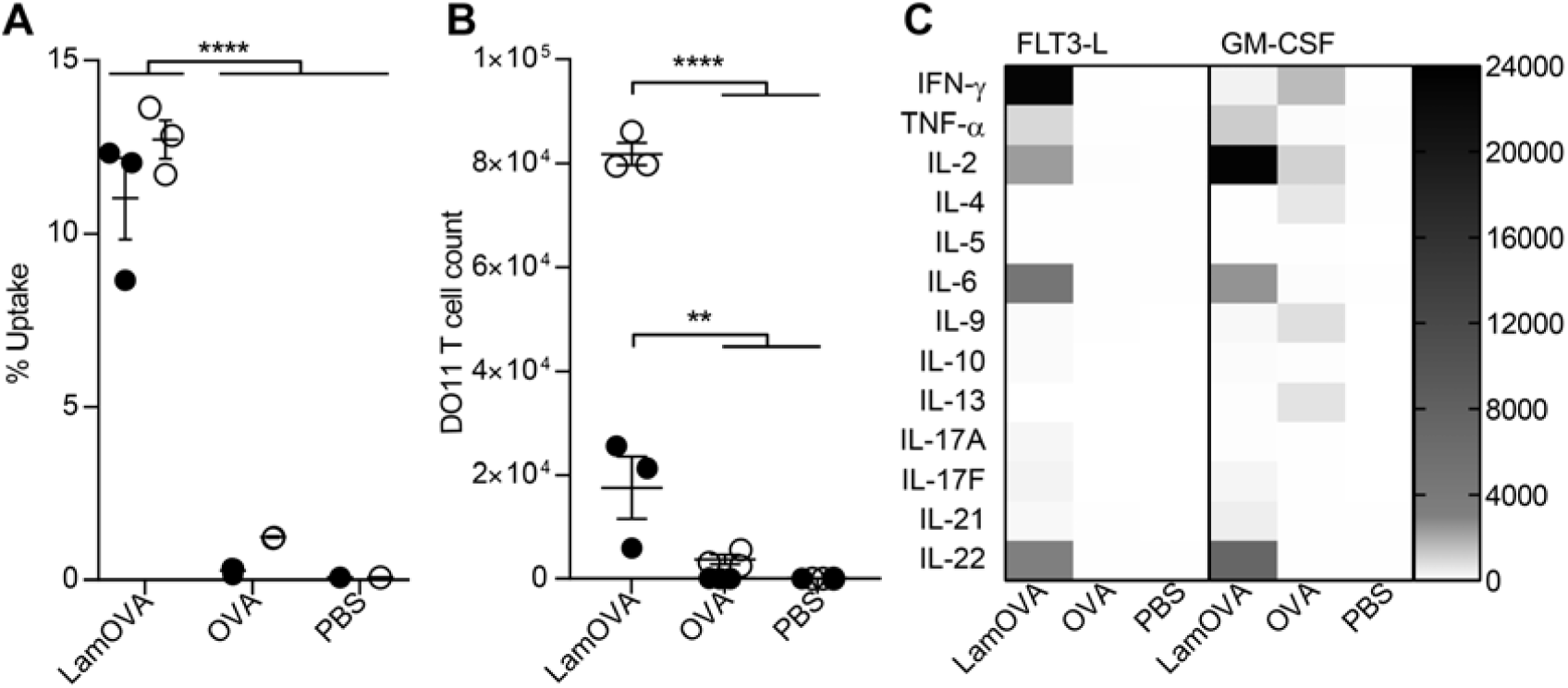
A) Uptake of pHrodo labeled OVA or LamOVA conjugates by GM-BMDCs (white circles) and FL-BMDCs (black dots) was analyzed by flow cytometry after 24h. B) Proliferation of OVA-specific DO11.10 T cells after 5d co-culture with BMDCs and OVA or LamOVA. Data are shown as number of live, proliferating DO11.10 cells. C) Cytokine concentration in BMDC-T cell co-culture supernatants in pg/mL. Means±SEM and individual technical replicates (n=3) are shown. Data were statistically analyzed by two-way ANOVA followed by Tukey’s post hoc test.

LamOVA-induced activation of BMDCs was associated with a robust, dose-dependent cytokine and chemokine response (Suppl. Fig. 3). Compared to OVA and laminarin, LamOVA significantly upregulated all measured cytokines and chemokines in FL-BMDCs (two-way RM ANOVA, followed by Tukey’s post hoc test). GM-BMDCs displayed a much higher basal activation status than FL-BMDCs as indicated by significantly higher levels of IL-1α, IL-1β, IFN-γ, IL-17A, IL-6, IL-27, CCL-3, CCL-4, CXCL10, CCL17, and CCL22 after 24h incubation without stimulus. CCL3, CCL4, CCL17, and CCL22 were especially elevated (37, 80, 138, and 101-fold) compared to FL-BMDCs. Despite this higher basal activation level, LamOVA significantly upregulated pro-inflammatory mediators (TNF-α, IFN-β, IL-6, IL-23, CCL3, CCL4, CCL5, CXCL10), and neutrophil attractants CXCL1 and CXCL5. In contrast, neither EndoFit™ OVA nor laminarin induced significant cytokine or chemokine secretion in either BMDC type.

### LamOVA activated BMDCs are potent inducers of T cell responses

Laminarin conjugation to OVA significantly increased uptake and induced activation of BMDCs and secretion of pro-inflammatory cytokines. These properties of LamOVA conjugates also resulted in enhanced stimulation of OVA-specific naïve T cells co-cultured with BMDCs. Both, FL- and GM-BMDCs incubated with LamOVA significantly activated T cell proliferation and cytokine secretion (Fig 2B and C). GM-BMDCs loaded with conjugates were more potent in inducing DO11.10 T cell proliferation than FL-BMDCs, which may be due to their higher basal activation status (suppl. Fig 2). This is also supported by the fact that only GM-BMDCs could induce activation of naïve T cells (Fig. 2B) and significant secretion of cytokines (IFN-γ and IL-9, Fig. 2C) in the presence of Endofit™ OVA. In both cell culture models, conjugates significantly induced IL-2, IL-6 and IL-22 production (P<0.0001, Fig 2C). Interestingly, LamOVA loaded FL- and GM-BMDCS displayed a different TH-polarizing potential as FL-BMDCs mainly generated TH1 cells (IFN-γ and TNF-α, P<0.0001 and P<0.001), whereas GM-BMDCs favored upregulation of IL-2, which correlated with the higher T cell proliferation rate.

### Epicutaneous immunization with LamOVA induces strong antibody responses

We have previously shown that epicutaneous immunization with mannan-conjugated allergen via laser-generated micropores (EPI) is considerably more immunogenic compared to immunization with unconjugated allergen ^24^. In a prophylactic immunization experiment (Fig. 3A), we could confirm that EPI with LamOVA induces ∼3-fold higher IgG1 antibody titers compared to OVA after two immunizations (Fig. 3B). This immune potentiating effect was dependent on the covalent linkage between OVA and laminarin since a mix of uncoupled OVA with laminarin did not enhance antibody responses. After a second booster immunization, all groups displayed similar antibody titers (Fig. 3C). IgG2a levels were close to the detection limit (data not shown).

**Fig. 3.**
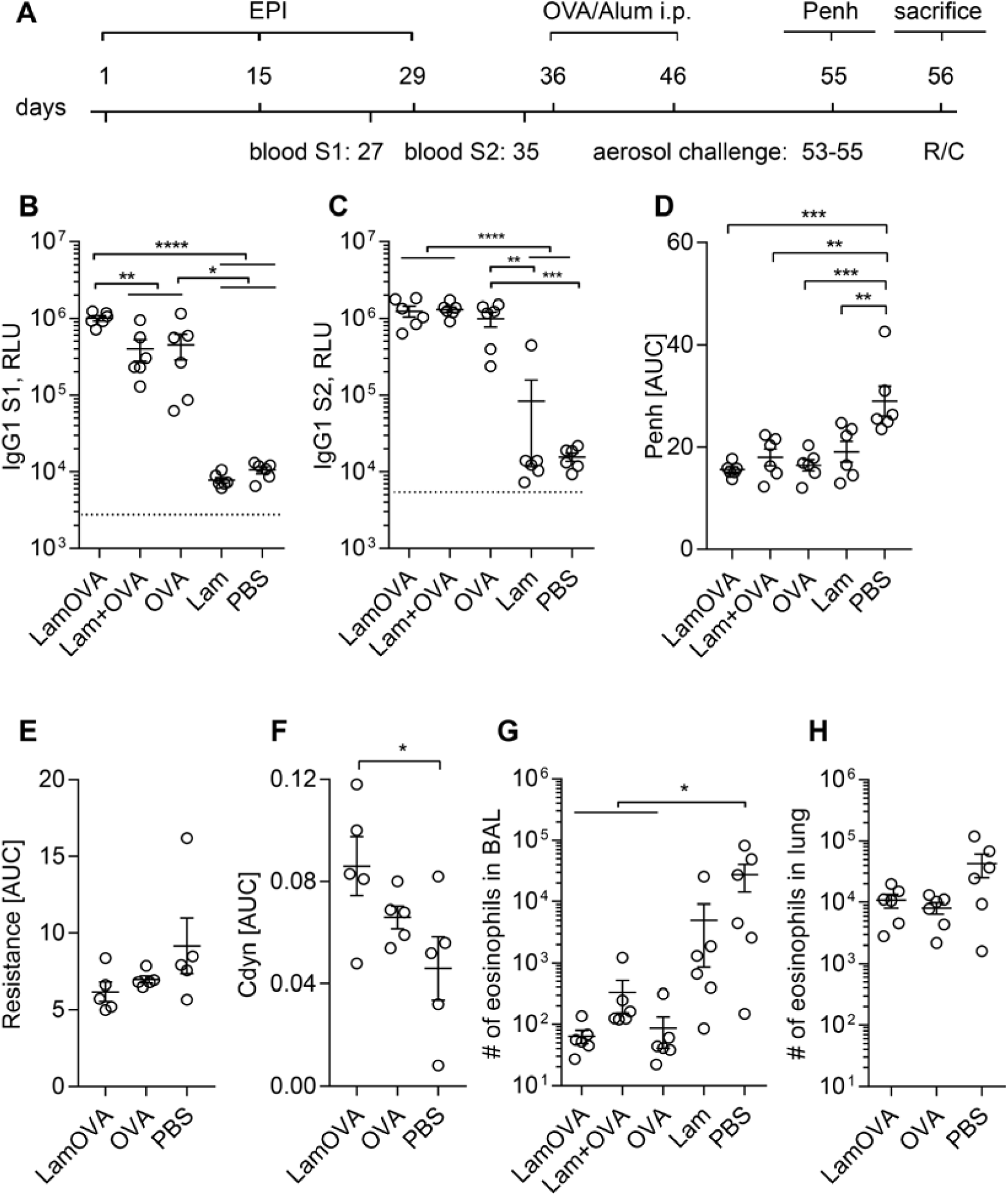
Prophylactic epicutaneous immunization via laser-generated micropores (EPI). A) Mice were immunized 3 times with LamOVA, OVA, a mix of OVA with laminarin (Lam+OVA), laminarin (Lam) or PBS followed by i.p. sensitization and aerosol challenge. Numbers indicate days. OVA-specific serum IgG1 was measured after two (B, S1) or three (C, S2) immunizations at a serum dilution of 1:10000. Data are shown as relative light units (RLU) of a luminometric ELISA (n=6). Background is indicated by dotted line. Airway hyperresponsiveness was assessed by WBP (D, n=6), as well as resistance (E, n=5) and compliance (F, n=5) measurements. Penh, resistance and compliance data are shown as area under the curve (AUC) of a methacholine challenge dose response curve.

### Preventive LamOVA immunizations reduce allergic lung inflammation and airway hyperresponsiveness

To investigate whether LamOVA-EPI would protect from allergic sensitization and lung inflammation, vaccinated mice were sensitized by two i.p. injection of Endofit™ OVA/alum followed by three aerosol challenges.

Lung function was measured by whole body plethysmography (WBP) under methacholine challenge. Mice vaccinated with LamOVA showed the lowest Penh values, significantly lower compared to the sham treated control group (Fig. 3D). The reduction of airway hyperresponsiveness (AHR) was confirmed in selected groups by invasive resistance (R) and compliance (Cdyn) measurement on the next day, confirming the WBP results (Fig. 3E and F).

After R/C measurement, the cellular composition and cytokine content of bronchoalveolar lavage fluid (BALF) were analysed. Preventive immunizations with OVA significantly reduced the number of eosinophils (Fig. 3G) in BALF, with no differences between the OVA, LamOVA, and Laminarin+OVA groups. Eosinophil counts in whole lung tissue confirmed BALF analysis for OVA and LamOVA (Fig. 3H). Eotaxin and IL-5 concentration in BALF was similar in all groups and did not correlate with eosinophil number in BAL and lung tissue (data not shown).

In summary, our data show that despite their hypoallergenicity, conjugates are more potent in the production of OVA-specific IgG antibodies and can prevent allergic sensitization and airway hyperresponsiveness. In a next step we therefore evaluated safety and potency of LamOVA conjugates in a therapeutic mouse model of allergic asthma.

### Epicutaneous immunotherapy with LamOVA induces blocking IgG and suppresses lung inflammation

In a therapeutic experiment, mice were sensitized by two i.p. injections of 10µg Endofit™ OVA/alum followed by intranasal (i.n.) OVA challenge to induce lung inflammation. Based on WBP data, mice were stratified into five treatment groups with similar means and distribution of Penh. RBL assay confirmed that all animals had OVA-specific IgE antibodies, however, treatment arms – by chance – showed slightly higher IgE levels before therapy compared to the control groups (suppl. Fig. 4). Mice were treated 8 times epicutaneously (EPIT) with either LamOVA or OVA. As a positive control, mice were treated by subcutaneous injections of Endofit™ OVA together with alum (OVA s.c.). Controls were sham treated with either PBS or laminarin (Fig 4A).

**Fig. 4.**
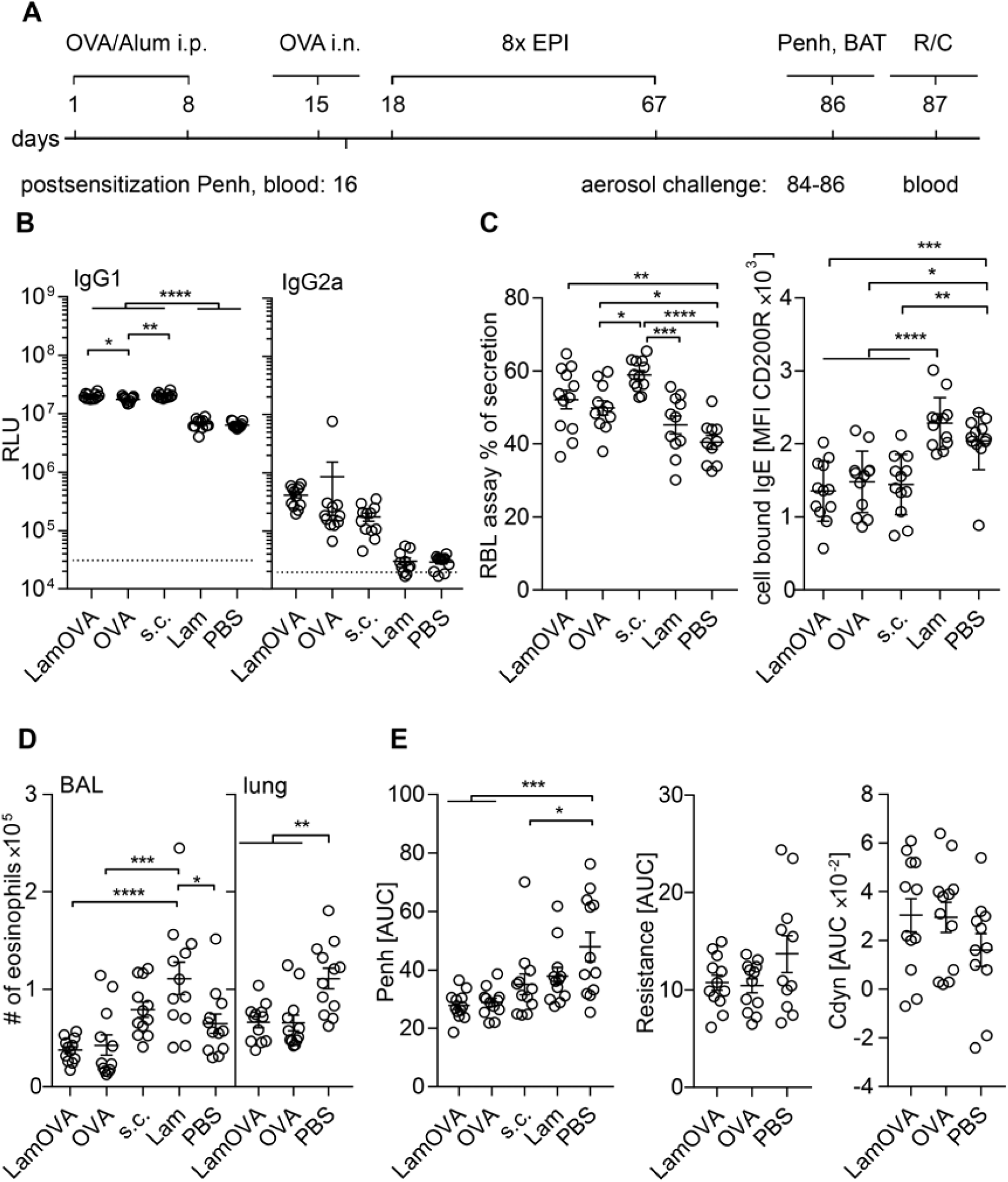
Therapeutic epicutaneous immunization via laser-generated micropores (EPIT). A) Mice were sensitized by two i.p. injection with OVA/alum and an intranasal challenge. After 8 treatments, either epicutaneously (LamOVA, OVA) or subcutaneously with OVA/alum (s.c.), mice were exposed 3 times to aerosolized OVA before sacrifice. Numbers indicated days. B) OVA-specific serum IgG1 and IgG2a at the timepoint of sacrifice were measured at a serum dilution of 1:10000 and 1:400, respectively. Data are shown as relative light units (RLU) of a luminometric ELISA (n=12). Background is indicated by dotted line. C) Serum and cell bound IgE were analyzed by RBL assay (serum dilution 1:150) and BAT, respectively. D) Number of eosinophils in BALF and lung tissue was analyzed by flow cytometry. E) Airway hyperresponsiveness was assessed by WBP as well as resistance and compliance (Cdyn) measurements. Penh, resistance and compliance data are shown as area under the curve (AUC) of a methacholine challenge dose response curve.

LamOVA was equally potent in inducing IgG1 antibody production compared to s.c. injections together with alum. In contrast, EPIT with unconjugated OVA elicited significantly less antibodies. Treatment with conjugates showed a trend towards increased production of IgG2a antibodies (Fig 4B).

Subcutaneous immunotherapy with OVA significantly increased serum IgE levels compared to the PBS treated control group. The IgE increase after EPIT with LamOVA and OVA was less prominent compared to SCIT with OVA/alum (Fig 4C). It is known that AIT can temporarily increase serum IgE levels. However, more relevant is the blocking of basophil and mast cell activation induced by cross linking of long-lived cell bound IgE. To test *ex vivo* blocking of basophil activation, blood samples were drawn 24h after the second exposure to aerosolized allergen and restimulated *in vitro* with OVA for 2h. Basophils from laminarin or PBS-treated control groups showed high activation levels as indicated by the strong expression of CD200R. In contrast, basophils from OVA-immunized groups showed significantly lower activation (Fig. 4C). To test, whether the reduced basophil activation was due to the presence of blocking IgG, blood samples from treated mice were either washed or left untreated before *in vitro* restimulation with OVA. In the absence of autologous serum, basophils from all groups showed similar activation (Suppl. Fig. 5A), indicating the presence of comparable amounts of OVA-specific IgE on the basophils. Thus, the IgG blocking capacity can be displayed as the ratio of basophils activation in the absence or presence of autologous serum. Correlating with the ELISA data, LamOVA EPIT group and OVA s.c. group showed the highest levels of blocking IgG (suppl. Fig. 5B).

EPIT groups displayed the lowest number of eosinophils in BALF and significantly lower numbers in lung tissue compared to sham treated mice (Fig. 4D). No significant differences in the BALF levels of IL-4, IL-5, IL-13, and eotaxin were found (data not shown). OVA treatment significantly improved lung function measured by WBP and again the EPIT groups showed the lowest Penh values (Fig. 4E). Though not statistically significant, a trend for reduced resistance and increased compliance in the EPIT groups was confirmed by invasive R/C measurement (Fig. 4E).

Restimulation of splenocytes with OVA (10ug/mL) induced increased expression of TH2 associated cytokines IL-4, IL-5, IL-10, and IL-13 but also IFN-γ, IL-2, IL-6, and IL-22 in the s.c. OVA group and surprisingly also in the group treated with laminarin alone. EPIT groups showed no such boost in cytokine responses and remained at similar levels compared to the PBS treated group (suppl. Fig. 6). This effect was also seen on the level of transcription factor expression Tbet, GATA3, and RORγT. No difference in the number of FoxP3+CD25+ Tregs was found in the spleens (suppl. Fig. 7). In summary, s.c. injection with OVA/alum as well as treatment with laminarin boosted mainly TH2, but also TH1 and TH22 responses, whereas EPIT with OVA or LamOVA induced no increase in cytokine responses compared to the untreated group.

### Conjugation of polysaccharide to OVA reduces local side effects in vivo

In mice, local side effects are dependent on the used allergen. While in previous studies with recombinant Phl p 5 ^25^ or depigmented house dust mite extract ^26^ no local side effects were observed, application of OVA to the skin of sensitized mice led to significant skin erythema and scab formation (Fig. 5A). We quantitated local skin reactions by calculating total erythema size and blob size (Fig 5B and C) and found that LamOVA immunizations induced significantly lower skin reactions, confirming the hypoallergenicity observed *in vitro*. The use of laser with PBS and laminarin alone did not induce any side effects, confirming the antigen-specificity of the response.

**Fig. 5.**
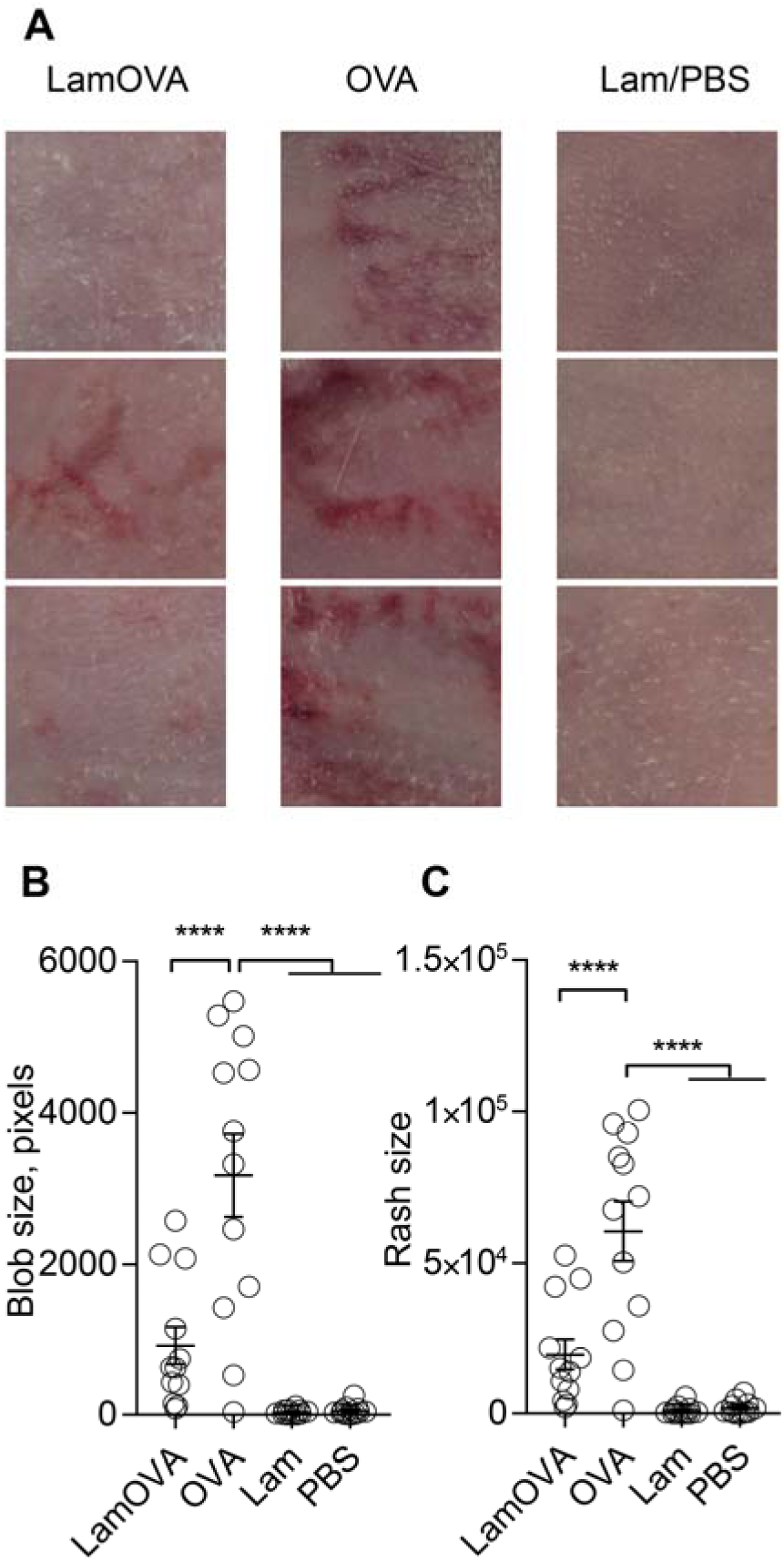
Skin erythema induced by EPIT A) 24 hours after the 4^th^ or 5^th^ treatment immunization sites were photographed and the sum of directly connected pixels of the segmented erythema (blob size, B) and the total erythema size (C) were quantitated (n=12). Panel A shows three representative animals from each group.

## Discussion

Allergen-specific immunotherapy (AIT) is the only disease-modifying approach for treatment of allergies. However, classical subcutaneous immunotherapy (SCIT) suffers from low patient compliance due to exhaustive treatment protocols and occurrence of local and systemic adverse reactions. Therefore, more efficient and safe therapeutic concepts are urgently needed. Subcutaneous injections deliver antigen into the hypodermis, an adipose tissue rich in blood vessels, but sparsely populated by APCs. In contrast, epidermis and dermis are rich in APCs and (epi)cutaneous immunotherapy may result in enhanced immunogenicity and efficacy. Delivery of allergen to superficial skin layers avoids contact with blood vessels, decreasing the risk of systemic side effects ^27, 28^. Upon epicutaneous application, antigen is taken up by Langerhans and dendritic cells, which migrate to the lymph nodes for presentation of antigenic peptides to T cells. In a peanut allergy model, EPIT with Viaskin^®^ occlusive patches led to formation of tolerogenic immune responses ^28^, demonstrating an important role of Langerhans cells in induction of regulatory T cells (Treg) in the context of skin immunization. Other APCs, which are able to generate Tregs, such as the CD11b+ cDC2 subset, have been shown to reside in the dermis ^29, 30^. Though epicutaneous allergen delivery with occlusive patches has been shown to effectively induce tolerance, this approach affords high amounts of allergen to be daily applied over several years ^31^ due to limited penetration of antigen through the outermost skin layer, the stratum corneum. In contrast, pre-treatment of the skin for disruption of this tight barrier can improve immunogenicity of EPIT. This approach has been introduced in Europe during the 1950s and has only recently proven its potency to reduce allergic symptoms in clinical trials ^2^. In this work, we utilized an infrared laser (P.L.E.A.S.E.^®^ Professional) device for controlled and highly reproducible barrier disruption. In addition to physically breaching the stratum corneum for facilitated antigen delivery directly to skin resident APCs ^16^, laser microporation creates a pro-inflammatory cytokine and chemokine milieu, which boosts immunogenicity ^32^.

Notably, Senti et al. reported that allergen application onto barrier disrupted skin may result in local and systemic side effects ^2^, thus highlighting the need for hypoallergenic, yet immunogenic formulations. Coupling of carbohydrates to allergens represents an elegant approach to reduce the IgE binding capacity of the resulting neoglycoconjugates, while simultaneously targeting and stimulating dendritic cells. We have reported that mannan-Bet v 1 conjugates (Bet-MN) are particularly hypoallergenic and no longer induce basophil activation *in vitro* ^15, 33^. In a similar approach, Sirvent et al. demonstrated hypoallergenicity of mannan allergoids using skin prick tests in allergic patients ^34^. In our current study, we found 6kD laminarin conjugates to be 5-fold less allergenic *in vitro* (Fig. 1B) and to induce significantly less local side effects in sensitized mice during therapy (Fig. 5). Coupling of allergens to β-glucan molecules with a higher molecular weight could potentially further decrease allergenicity as we have previously seen a >1000-fold reduction of IgE crosslinking capacity of 30-40kD mannan conjugates ^15^. However, a careful balance of protein/carbohydrate ratio has to be maintained, as a higher polysaccharide content in conjugates potentially decreases IgG production (unpublished observation).

Due to the low immunogenicity of existing allergy vaccines, AIT requires repetitive administrations of high allergen doses. More potent and controlled APC activation may be achieved via specific targeting of surface receptors ^6^, e.g. C-type lectin receptors ^35^. Here, we used laminarin to target dectin-1, which is expressed on various cell types including CD11b+ dermal DCs and Langerhans cells in mice ^36^. We ^15, 33^ and others ^34, 37^ have previously reported enhanced uptake of neoglycoconjugates compared to unconjugated protein. In contrast to Sirvent et al., who observed that oxidation of mannan impaired uptake of conjugates ^34^, we have shown that mild oxidation of laminarin for allergen coupling does not influence its biological activity, as confirmed by its binding to dectin-1 (Fig. 1A) and activation of BMDCs (suppl. Fig. 2 and 3). We previously made similar observations for mannan ^15^ and other carbohydrates ^33^. We employed two different in vitro models for studying antigen uptake, DC activation, and stimulation of naïve T cells, i.e. Flt3L derived BMDCs and GM-CSF derived BMDCs. While FL-BMDCs more closely represent steady state naïve DCs, GM-BMDCs rather resemble monocyte-derived inflammatory DCs (and macrophages) ^20, 21^. Both models are relevant for skin vaccine evaluation as they represent both, the early phase of the immune response, when naïve DCs first encounter an antigen, and the late phase response, when local inflammation (induced by adjuvants or laser microporation) attracts inflammatory DCs to the vaccination site. Indeed, GM-BMDCs displayed a higher basal activation status in the absence of stimuli as indicated by increased levels of CD86 and dectin-1 expression (suppl. Fig 2) and enhanced secretion of cytokines and chemokines (suppl. Fig. 3) compared to FL-BMDCs. Laminarin is a β-1,3-glucan and, depending on the mode of uptake, can have pro- or anti-inflammatory properties. For example, low molecular weight laminarins do not stimulate APCs ^38^, as dectin-1 activation occurs only after clustering of several units of the receptor ^39^. Hence, receptor oligomerization requires a specific size of β-glucan complexes ^38^. Here, we demonstrate that (dialyzed) 6kDa laminarin does not activate BMDCs in vitro, whereas laminarin-OVA complexes are very potent in stimulating GM-BMDCs as well as FL-BMDCs. It has been shown that receptor internalization attenuates pro-inflammatory signaling, whereas large particles such as zymosan cannot be internalized and induce strong pro-inflammatory immune responses ^40^. Corroborating findings from Xie et al. ^37^, our data indicate that LamOVA is taken up by receptor-mediated phagocytosis as indicated by downregulation of surface dectin-1 expression (suppl. Fig. 2) and transport into the endosome (Fig. 2A). Yet, in contrast to data from others ^40^, we clearly observed the induction of inflammatory pathways in LamOVA stimulated BMDCs. In line with the BMDC activation status, only GM-BMDCs could activate naïve OVA-specific T cells without additional stimuli (Fig. 2B and C). In contrast, BMDCs pulsed with LamOVA were potent activators of naïve T cell proliferation and cytokine secretion. As components of fungal cell walls, β-1,3-glucans are known to trigger Th1/Th17 adaptive immunity ^36, 41^. This could be confirmed in T cells stimulated with LamOVA loaded FL-BMDCs, which secreted high levels of IFN-γ, TNF-α, and IL-22, and low levels of IL-17 (Fig. 2C). Interestingly, although GM-DCs induced stronger T cell proliferation and IL-2 secretion, they were less potent in inducing TH1 cytokine secretion, which also correlates with their lower levels of IL-12 secretion upon LamOVA stimulation compared to FL-BMDCs (suppl. Fig. 3).

We have recently demonstrated that application of allergens to laser-microporated skin can induce humoral immune responses ^16, 26, 42^ and that conjugation to mannan potentiates antibody responses ^15^. As shown previously for mannan, laminarin also boosted IgG1 responses 3-fold after EPI and covalent linkage of the carbohydrate to OVA was essential for this effect (Fig. 3B). A similar observation was made by Xie et al. after s.c. immunization, however they also included poly I:C as an adjuvant ^37^. In a prophylactic setting, EPI with OVA and LamOVA significantly suppressed induction of allergen-induced lung inflammation. Although statistically not significant, LamOVA vaccinated animals showed lower lung resistance (Fig. 3E) and higher dynamic compliance (Fig. 3F) compared to OVA immunized mice. Thus, despite priming TH1/TH17 responses in vitro, LamOVA had no detrimental inflammatory effects in a prophylactic allergy model in vivo.

We have previously shown in a therapeutic mouse model of allergic lung inflammation that laser-mediated EPIT with Phl p 5 or house dust mite extract can successfully reduce lung inflammation and improve lung function. Therapeutic efficacy was associated with induction of high levels of blocking IgG and a general downregulation of cytokine responses ^25, 26^. In the current study, we show for the first time the therapeutic efficacy of a hypoallergenic laminarin-allergen conjugate. While EPIT with OVA induced severe local side effects (Fig. 5), replicating findings from clinical trials with grass pollen extract ^2^, LamOVA treated mice showed significantly ameliorated skin erythema. At the same time, conjugation to laminarin significantly boosted IgG1 to the same levels as achieved by s.c. injections together with alum (Fig. 4A), while inducing lower levels of therapy-associated IgE (Fig. 4C). Furthermore, both EPIT groups showed lower levels of lung inflammation compared to s.c. treated mice (Fig. 4D and E). Although LamOVA induced inflammation and TH1/TH17 polarization in vitro, EPIT with LamOVA did not result in enhanced inflammatory cytokine secretion in splenocytes from treated animals (suppl. Fig. 6). In contrast, SCIT with OVA boosted TH2 responses, an early therapy effect that is well known in the clinics ^43^. Though we did not find increased numbers of FoxP3+CD25+ regulatory T cells in the spleen after treatment, EPIT significantly reduced the number of activated T cells compared to SCIT. Taken together, this confirms our previous findings that EPIT via laser-generated micropores can downregulate established allergic responses and mainly results in the induction of blocking IgG and a general suppression of cytokines ^25, 26^ rather than immune deviation towards TH1 or TH17, even in the presence of an immunostimulatory dectin-1 agonist. While the first clinical studies of SCIT with mannan allergoids are currently under way (NCT02654223, NCT02661854), we believe that combining a C-type lectin receptor based DC targeting approach with a method to deliver a hypoallergenic vaccine to upper skin layers rich in APCs is the key to patient friendly, safe and effective immunotherapy. Our current proof of concept study demonstrates that laser-facilitated EPIT with β-glucan-allergen conjugates represents one such feasible approach.

## Acknowledgements

This work was supported by the Austrian Science Fund (FWF; grant no. W 1213).

