## Supplementary Material for "Laser-facilitated epicutaneous immunotherapy with hypoallergenic beta-glucan neoglycoconjugates suppresses lung inflammation and avoids local side effects in a mouse model of allergic asthma"

### Supplementary Methods

Generation of bone marrow-derived dendritic cells

Bone marrow was harvested from femur and tibia of BALB/c mice. For generation of FL-BMDCs, 3 mL of a 2.5x10^6^ cells/mL bone marrow cell suspension were seeded into non-cell culture treated 6-well plates and stimulated with 200ng/mL human Flt3-L for 9 days at 37°C, 5% CO_2_ and 95% humidity. Between days 4 and 6, 1.5 mL of culture medium (RPMI/1640 supplemented with 2.5 mM HEPES, 100 IU/mL penicillin, 100 mg/mL streptomycin, 2 mM L-glutamine, 10% fetal calf serum) were added to each well. For generation of GM-BMDCs, 10 mL of a 2x10^5^ cells/mL bone marrow cell suspension were plated into non-cell culture treated petri dishes and incubated in the presence of murine GM-CSF for 8 days. On day 3, 10 mL of GM-CSF containing culture medium (RPMI/1640 supplemented with 20 ng/mL GM-CSF, 100 IU/mL penicillin, 100 mg/mL streptomycin, 2 mM L/Glutamine, 10% FCS and 50 µM 2-mercaptoethanol) was added to each well, and on day 5, 10 mL medium was replaced.

Conjugate uptake, BMDC activation and T cell proliferation

To assess uptake of glycoconjugates by BMDCs, their activation, and their capacity to induce T cell proliferation, FL-BMDCs and GM-BMDCs harvested on days 9 and 8, respectively were used.

For uptake analysis, OVA and LamOVA were labeled with Invitrogen pHrodo™ iFL Green STP Ester (ThermoFisher Scientific) according to the manufacturer’s protocol. Briefly, pHrodo was resuspended in DMSO to a concentration of 10 mM. For labeling, 50 µL OVA or glycoconjugates were mixed with 5 µL of a 1M NaHCO_3_ solution (pH 8.3), added to the dye and incubated at RT for 60 min in the dark. The coupling reaction was stopped by addition of 1/10 volume of 1 M Tris pH8 followed by another incubation for 60 min at RT in the dark. Unreacted dye was removed from the samples by running over an Illustra NAP-5 column (GE healthcare). Labeled OVA and conjugates were analyzed for their fluorescence intensity at an excitation wavelength of 488nm and an emission wavelength scan between 520 and 560nm. Exact protein concentrations were determined by amino-acid analysis.

1.5x10^5^ cells were stimulated overnight at 37°C, 5% CO_2_, 95% humidity with 5µg of labeled conjugates or OVA, respectively. Incubation with PBS served as negative control. To analyze CD86 and Dectin-1 expression, 1.5x10^5^ cells were stimulated with 5, 25 or 50 µg/mL OVA or an equivalent amount of LamOVA, 12.5, 62.5 or 125 µg/mL laminarin, 1µg/mL LPS (positive control), or PBS (negative control), respectively, overnight at 37°C, 5% CO_2_, 95% humidity. Cells were analyzed by flow cytometry on a Cytoflex S flow cytometer (Beckman Coulter). See suppl. Fig. 8 for gating strategy.

For T cell – BMDC co-cultures, OVA-specific T cells harvested from BALB/c DO11.10 mice were labeled with 1 µM fluorescent dye carboxyfluorescein diacetate succinimidyl ester (CFSE) in PBS. Briefly, lymph node and spleen cell suspensions were incubated for 10 minutes at 37°C in a water bath with CFSE. The reaction was stopped by addition of 10% FCS. After washing, the cells were resuspended in T cell medium (RPMI-1640 supplemented with 25 mM HEPES, 100 IU/mL penicillin, 100 mg/mL streptomycin, 2 mM L-glutamine, 10% FCS) and stained with anti-CD62L – APC-eFluor780 and anti-CD4 – eFluor450 for sorting of naïve T cells on a BD FACS Aria^TM^ III cell sorter (BD Biosciences). Naïve DO11.10 T cells (1x10^5^ cells/mL) and BMDCs (0.33x10^5^ cells/mL) were co-cultured in T cell medium in the presence of 50 µg/mL OVA or an equivalent amount of conjugates or medium for 5 days at 37°C, 5% CO_2_, 95% humidity. To determine the percentage of proliferating cells, staining with anti-CD4-PerCP-Cy5.5 and fixable viability dye APC-eFluor 780 was performed and cells were analyzed by flow cytometry. Culture supernatants were used for cytokine secretion analysis by LEGENDPlex assay according to manufacturer’s protocol.

BAL

Bronchoalveolar lavages were performed as described previously ^1^. Following staining with anti-CD8-FITC, anti-SiglecF-PE, anti-CD45-PerCp-Cy5.5, anti-CD4‐BV421, anti-CD19‐PE/Cy7, and anti-Ly6G/Ly6C (Gr1)‐APC, cells were analyzed by flow cytometry. Leucocytes were gated on a SSC vs. CD45 plot followed by doublets exclusion. Neutrophils were determined as Gr1^high^ Siglec F^low^, and Gr1^low^ Siglec F^high^ cells were further divided into eosinophils and monocytes based on autofluorescence in the BV510 channel. Gr1^neg^ Siglec F^neg^ lymphocytes were further separated into CD4 and CD8 T cells. See suppl. Fig. 9 for gating strategy.

Lung digestion

After transcardial perfusion, the right lung lobe was cut into 1x1mm pieces in digestion buffer containing Liberase™ (0.25 mg/mL, Roche), Hyaluronidase (1 mg/mL, Sigma), DNAse I (0.03 mg/mL, Sigma) and Collagenase XI (0.25 mg/mL, Sigma) in RPMI-1640 and incubated on a rotary shaker (300rpm, 30min, 37°C). Thereafter, the cell suspension without visible clumps was placed on ice and the remaining tissue was passed through a 21G needle and combined with 2 mL of fresh digestion buffer. The suspension was incubated for additional 30 min under the same conditions. The cells were pooled and EDTA was added to a final concentration of 10mM. The sample was filtered through a 100µm cell strainer (Greiner Bio One) and labeled (15min, 4°C) with biotin anti-mouse CD45.2 antibody diluted 1:50 in FACS buffer (PBS, 1% BSA, 2mM EDTA) and sorted using BD IMag™ Streptavidin Particles Plus – DM (BD Biosciences). Prior to staining, non-specific binding sites were blocked by resuspending cells in 20µL hybridoma supernatant containing anti-CD16/CD32 for 5 min at 4°C. Finally, the cell pellet was stained using the following anti-mouse antibodies: CD11b- PerCp/Cy5.5, CD45-eFluor506, GR-1-AlexaFluor700, Siglec F-Superbright 600, CD24-PE/Dazzle 594, MHC II-FITC. After incubation for 30 min at 4°C, cells were washed tree times. Finally, the pellet was resuspended in FACS buffer and cells were analyzed on a Cytoflex S flow cytometer (Beckman Coulter). Samples were analyzed according to the gating strategy published by Misharin et al. ^2^ (Suppl. Fig. 10).

Basohphil activation test (BAT)

Cell-bound IgE was analyzed *ex vivo* by basophil activation test. Whole blood was drawn from vena saphena the day after the second aerosol challenge. To prevent coagulation, 10% v/v Li-Heparin (10mg/mL in DPBS) was added. 30µL of heparinized blood was either used untouched or washed three times with 200µL of RPMI (260g, RT, 5min) to remove serum containing blocking IgGs. 30µL of RPMI with or without 2ng/mL OVA was added to the samples and incubated for 2h at 37°C, 5% CO2, 95% humidity. Subsequently, samples were put on ice and all further steps were performed with ice-cold solutions at 4°C. Samples were washed with 120µL FACS buffer and pelleted (5min centrifugation at 260 g). Cells were stained in 30µL of FACS buffer containing anti-mouse IgE-FITC, anti-CD4-PerCp-Cy5.5, anti-CD19-PE/Cy7, and anti-CD200R-APC for 20 min on ice, washed with 200µL FACS buffer, and the pellets were resuspended in red blood cell lysis buffer (eBioscience) and incubated for 5min at RT followed by two washing steps with FACS buffer. Finally, cells were resuspended in 50µL FACS buffer and analyzed by flow cytometry on a FACS Canto II flow cytometer (BD Biosciences). Basophils were gated as IgE^high^ CD19^neg^ CD4^neg^ and the basophil activation status was assessed as median fluorescence intensity of CD200R.

Flow Cytometry of restimulated splenocytes

Cell pellets from restimulated cells were centrifuged for 5min at 400g at 4°C. After one wash with 150µL DPBS, pellets were resuspended in 20 µL hybridoma supernatant containing anti-CD16/32 to block Fc-receptors. After 5min incubation at 4°C, 20 µL of extracellular staining mix was added directly to the samples. Extracellular staining mix contained fixable viability dye eFluor 506, anti-CD44-BV650, anti-CD25-Super Bright 600 anti-CD4-APC-eFluor780 diluted in DPBS. Plates were briefly mixed on an oscillating shaker and incubated for 30 min at 4°C in the dark. Subsequently, 100µL of ice-cold FACS buffer were added, and cells were pelleted (400g, 4 min, 4°C). Cell pellets were resuspended in 150 µL ice-cold Fix/Perm buffer (Foxp3/Transcription Factor Staining Buffer Kit, Tonbo Biosciences) and incubated 60min at 4°C in the dark. After centrifugation (400g, 4min, 4°C), pellets were washed twice in 100 µL Perm buffer from the same kit. Pellets were resuspended in 30 µL of Perm buffer containing anti-FoxP3-APC and GATA3-BV421. Samples were incubated for 30 min at RT in the dark. After one more wash with 100 µL Perm buffer, cell pellets were resuspended in 80 µL FACS buffer and analyzed on a Cytoflex S flow cytometer (Beckman Coulter). See suppl. Fig. 11 for gating strategy.

### Supplementary Tables

| Marker | Fluorophore | Final Dilution | Clone | Manufacturer |
| --- | --- | --- | --- | --- |
| **BMDC stimulation, BMDC-DO11 T cell co-culture** | | | | |
| CD11b | PerCP-Cy5.5 | 1:500 | M1/70 | BioLegend |
| CD11c | BV421 | 1:100 | N418 | BioLegend |
| CD86 | BV605 | 1:200 | GL-1 | BioLegend |
| Dectin-1 | PE | 1:100 | RH1 | BioLegend |
| Fixable viability dye | APC-eFluor780 | 1:3000 |  | Invitrogen |
| **Lung digestion** | | | | |
| CD11b | PerCp-Cy5.5 | 1:200 | M1/70 | BioLegend |
| CD24 | PE/Dazzle 594 | 1:600 | M1/69 | Biolegend |
| CD45 | eFluor 506 | 1:100 | 30-F11 | Invitrogen |
| GR-1 | Alexa Fluor 700 | 1:200 | RB6-8C5 | Biolegend |
| MHC II | FITC | 1:200 | M5/114.15.2 | BioLegend |
| Siglec F | Superbright 600 | 1:200 | 1RNM44N | Invitrogen |
| BAL | | | | |
| CD4 | BV421 | 1:200 | RM4-5 | Invitrogen |
| CD45 | PerCP-Cy5.5 | 1:400 | A20 | Invitrogen |
| GR-1 | APC | 1:200 | RB6-8C5 | Invitrogen |
| Siglec F | PE | 1:200 | E50-2440 | BD |
| CD8 | FITC | 1:100 | 53-6.7 | Invitrogen |
| CD19 | PE-Cy7 | 1:100 | 1D3 | Invitrogen |
| **Lymphocyte restimulation** | | | | |
| CD4 | APC-eFluor780 | 1:400 | RM4-5 | Invitrogen |
| CD25 | Superbright 600 | 1:100 | PC61.5 | Invitrogen |
| CD44 | BV650 | 1:50 | IM7 | Biolegend |
| FoxP3 | APC | 1:100 | FJK-16s | Invitrogen |
| GATA3 | BV421 | 1:100 | 16E10A23 | BioLegend |
| Fixable viability dye | eFluor 506 | 1:1000 |  | Invitrogen |
| **BAT** | | | | |
| CD4 | PerCp-Cy5.5 | 1:200 | GK1.5 | BioLegend |
| CD19 | PE-Cy7 | 1:200 | eBio1D3 (1D3) | eBioscience |
| IgE | FITC | 1:200 | RME-1 | BioLegend |
| CD200R | APC | 1:200 | OX110 | Invitrogen |

**Supplementary Table 1. Antibodies used in the study.**

### Supplementary Figures

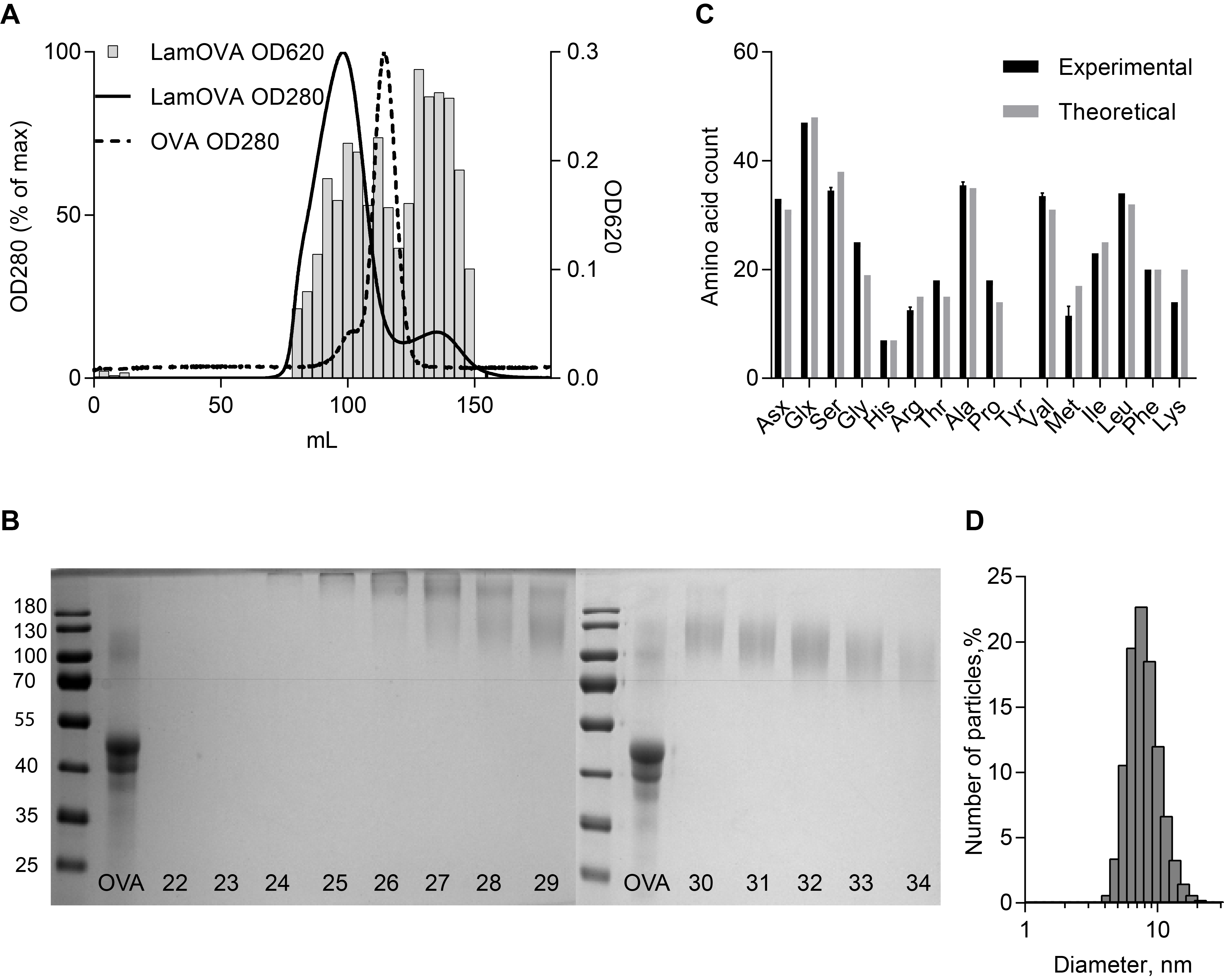

**Supplementary Figure 1. Characterization of laminarin OVA conjugates.** A) SEC chromatograms for LamOVA and OVA at 280nm. Laminarin concentration in LamOVA fractions was determined by anthrone method (620nm). B) OVA and LamOVA fractions 22-34 were analyzed by 10% SDS-PAGE and coomassie staining. Fractions 24-34 (elution between 70 and 100mL) migrated above 70kDa and were pooled for further analysis. C) Amino acid analysis. After reductive amination, only 14 of 20 lysine residues were detected. D) LamOVA size distribution (number-weighted) of particles as measured by dynamic light scattering. Mean hydrodynamic diameter was 8.27nm.

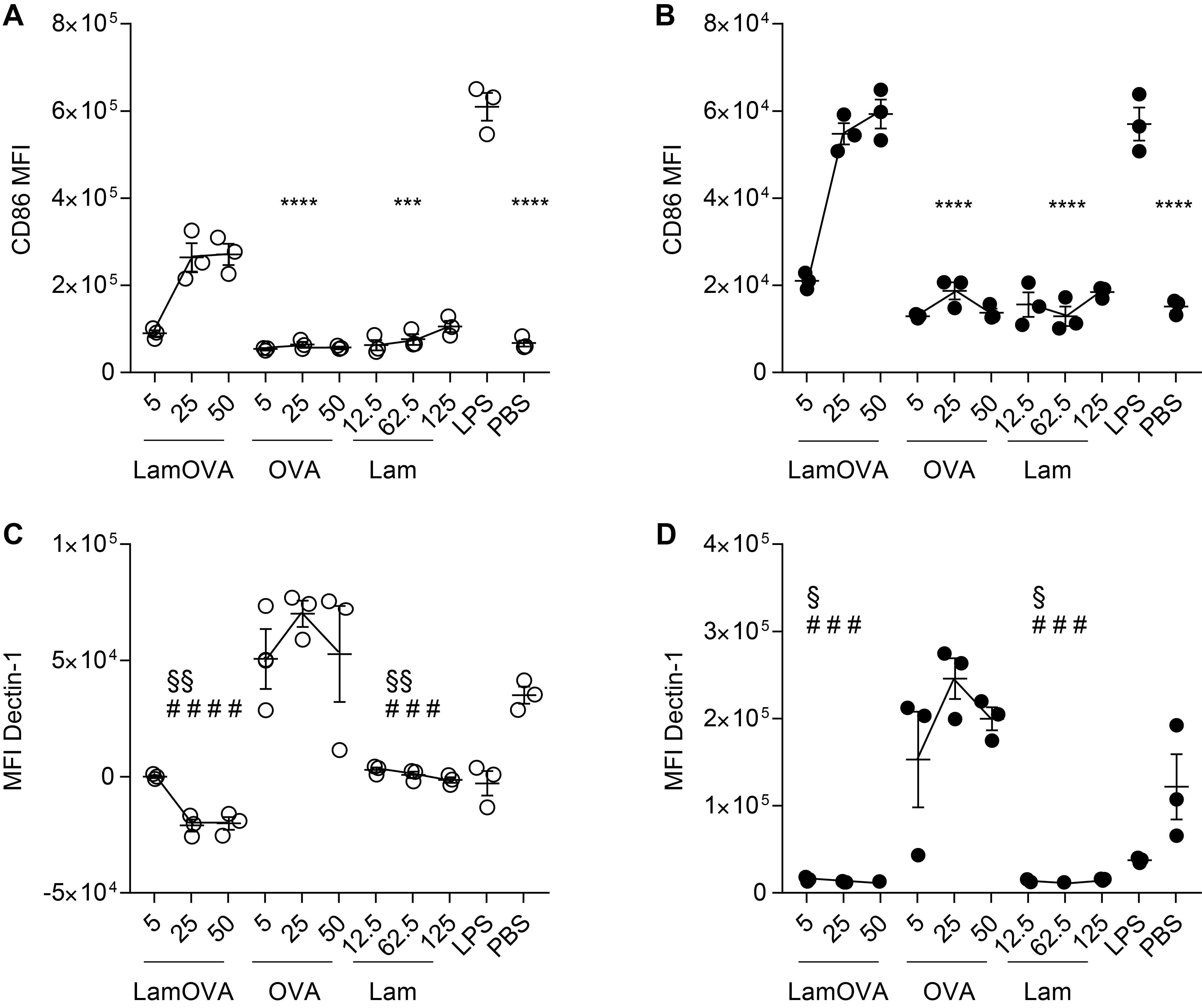

**Supplementary Figure 2. CD86 and dectin-1 expression on stimulated BMDCs**. GM- and FL-BMDCs were stimulated with LamOVA, OVA, or PBS for 24 hours. Protein concentrations for OVA and LamOVA were 5, 25, and 50µg/mL. Laminarin was used at equivalent doses as those present in LamOVA (12.5, 25, 62.5µg/mL). LPS (1µg/mL) served as positive control. Data are show as median fluorescence intensity (MFI) of CD86 (A and B) and dectin-1 (C and D) on GM-BMDCs (A and C, white circles) and FL-BMDCs (B and D, black dots). Data are shown as means±SEM and individual technical replicates (n=3). Statistical significance vs. LamOVA (*), vs. OVA (#), and vs. PBS (§) was analyzed by two-way RM ANOVA followed by Tukey’s multiple comparisons test.

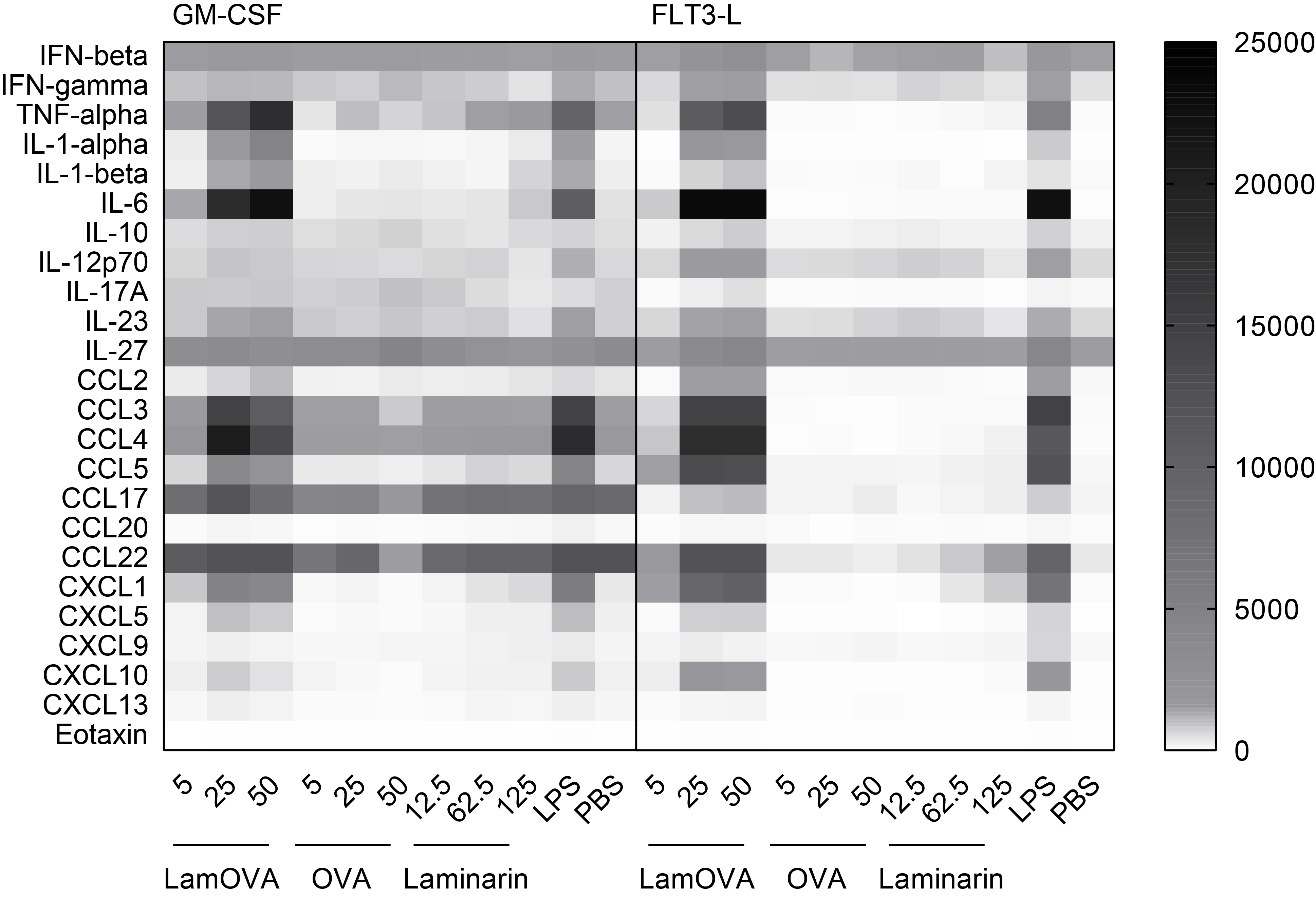

**Supplementary Figure 3. Cytokine and chemokine secretion from stimulated BMDCs.** GM- and FL-BMDCs were incubated 24h in the presence or absence of increasing doses of OVA, LamOVA or laminarin (shown in µg/mL). Laminarin was used at the same doses as present in LamOVA conjugates. 1 µg/mL LPS served as positive control. Data are shown as pg/mL.

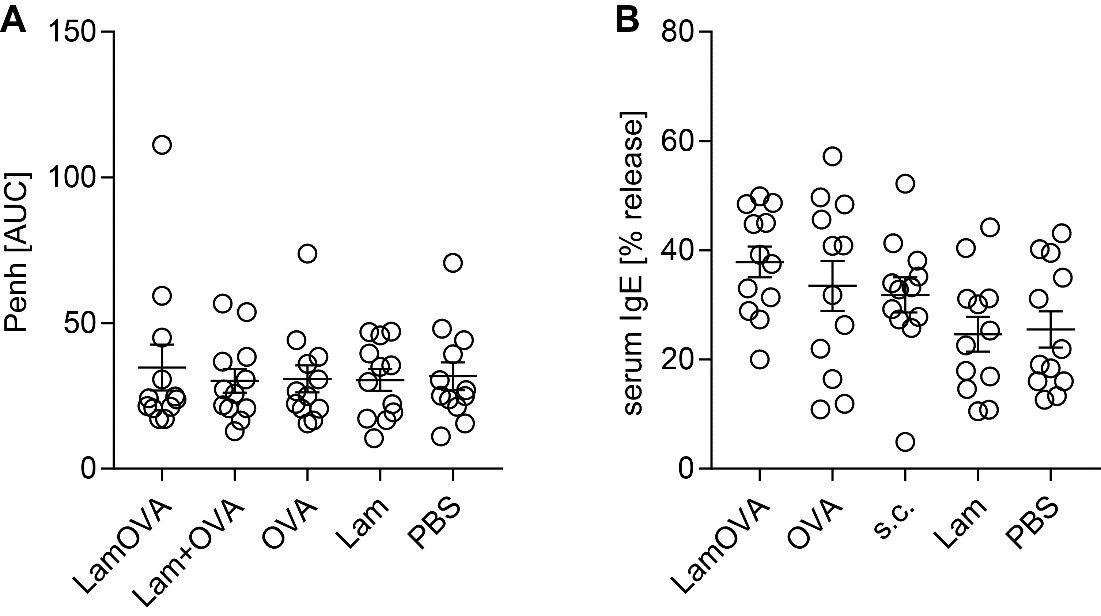

**Supplementary Figure 4. Group stratification of sensitized mice before treatment**. BALB/c mice were sensitized by two i.p. injections of OVA followed by two i.n challenges. A) Before treatment, airway hyperresponsiveness was assessed by WBP and mice were stratified into treatment groups with similar means and distribution of Penh. Data are shown as area under the curve (AUC) of a methacholine challenge dose response curve. B) Presence of OVA-specific IgE before treatment was confirmed by RBL assay (serum dilution 1:100). Data are shown as percentage of total beta-hexosaminidase release.

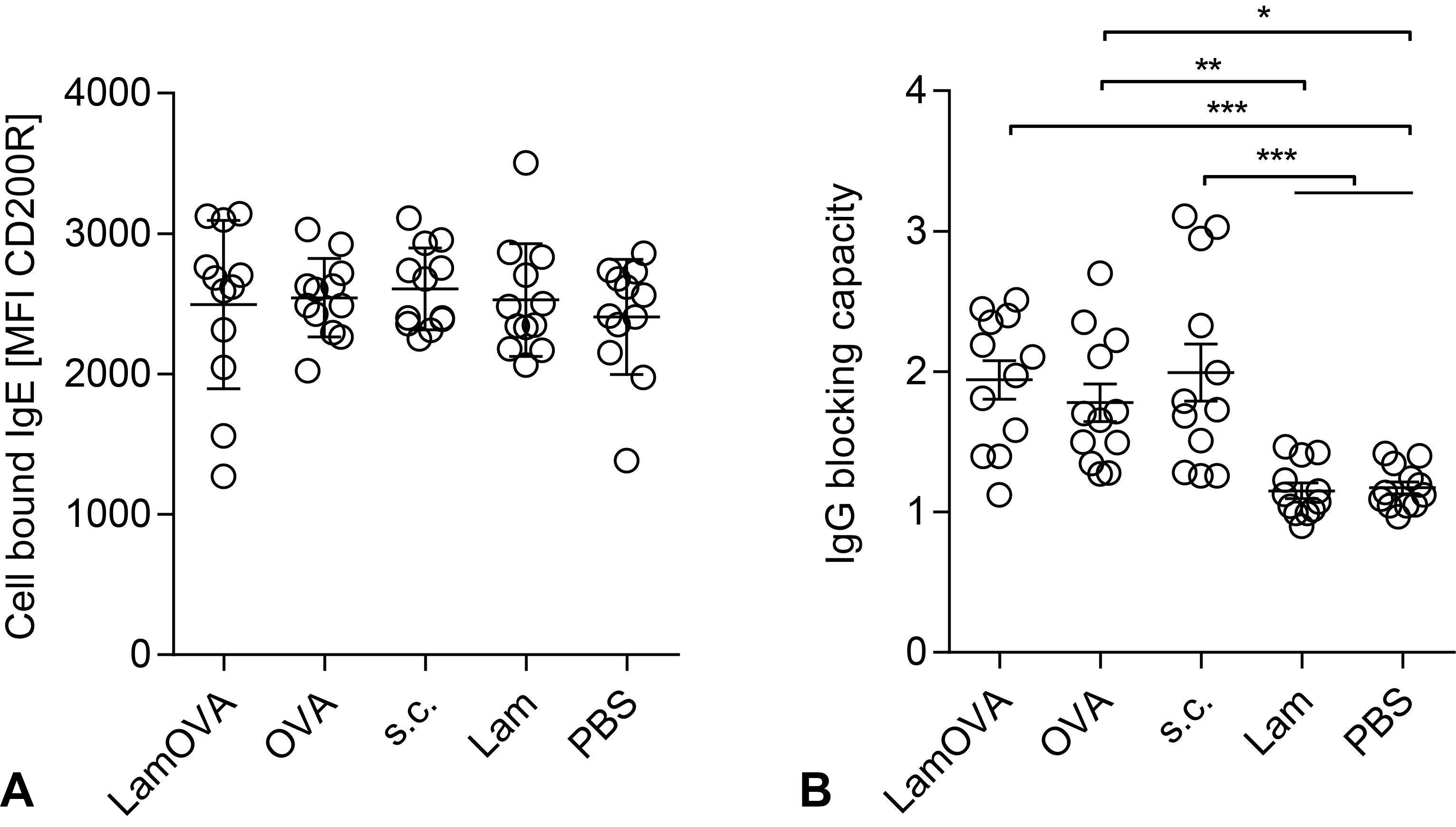

**Supplementary Figure 5. Cell bound IgE and IgG blocking capacity after treatment and challenge**. A) After the second aerosol challenge, blood was drawn and serum was washed away. After 2h *in vitro* restimulation with OVA, basophil activation status was assessed by flow cytometry as upregulation of CD200R. MFI = median fluorescence intensity. B) IgG blocking capacity is shown as the ratio of CD200R expression in basophils in the absence and presence of autologous serum. Data are shown as means±SEM and individual animals (n=12).

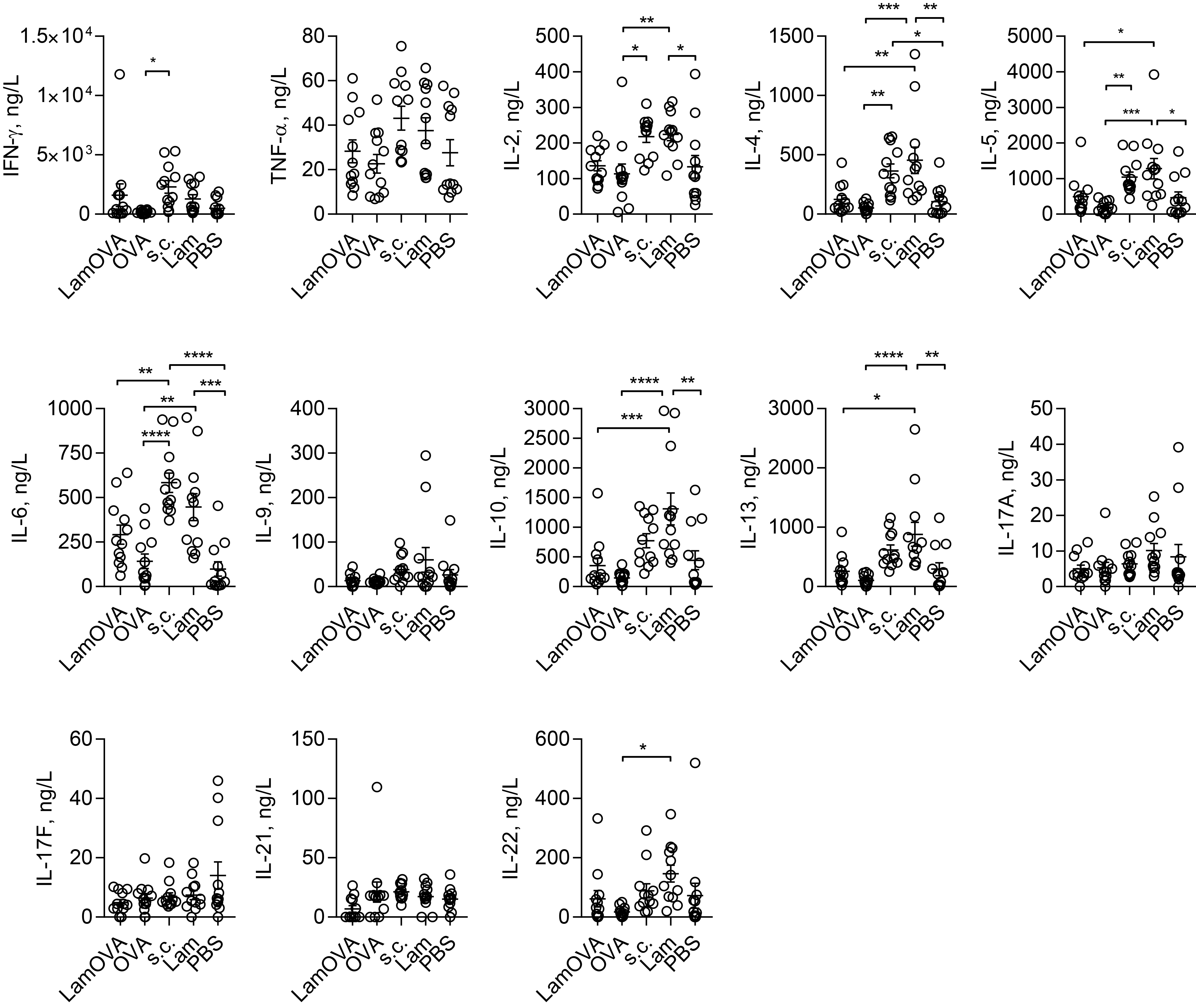

**Supplementary Figure 6. Cytokine secretion of splenocytes restimulated with OVA for 3 days.** Concentrations of different cytokines in cell culture supernatants was analyzed by flow cytometry using LEGENDplex assay. Data are shown as means±SEM and individual animals (n=12).

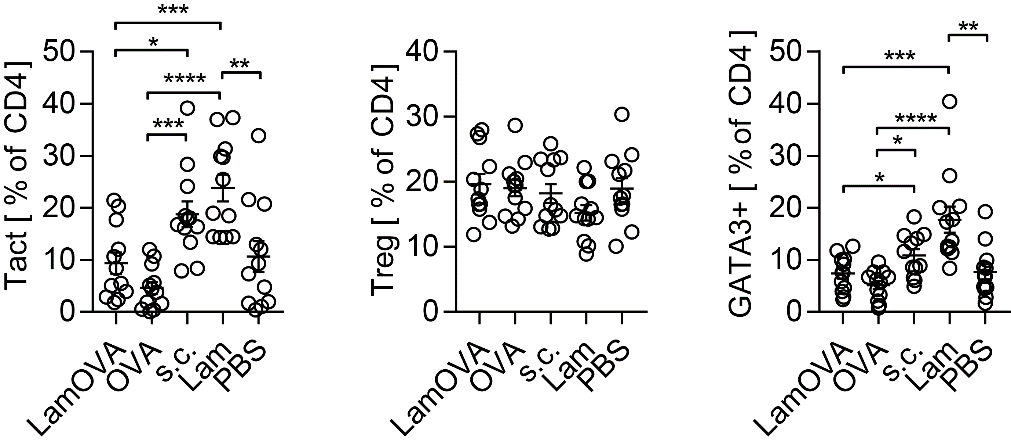

**Supplementary Figure 7. Percentage of activated (Tact), regulatory (Treg) and GATA3+ T cells.** Splenocytes were restimulated with OVA for 3 days and the percentage of Tact, Treg, and GATA3+ cell was assessed by flow cytometry. Data are shown as means±SEM and individual data points (n=12).

A	B

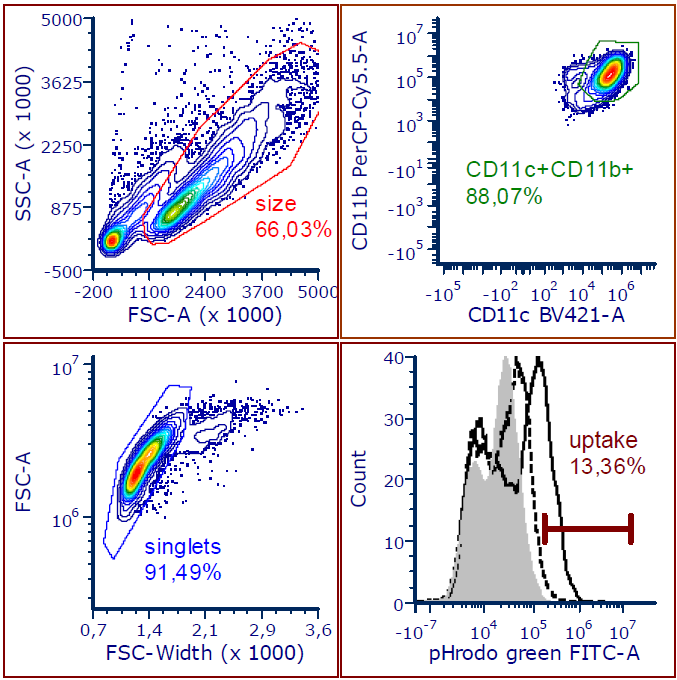

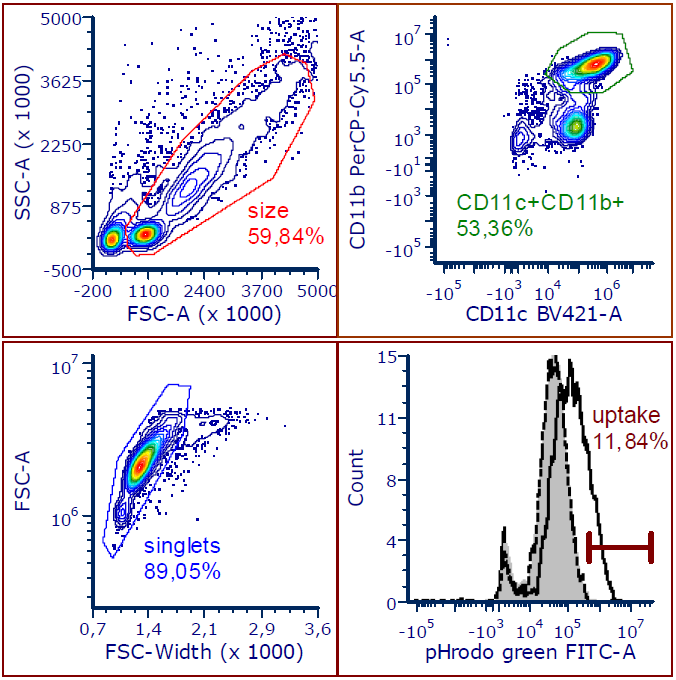

**Supplementary Figure 8. Gating strategy for BMDCs**. GM-BMDC (A) or FL-BMDCs (B) were first gated according to size to remove debris. Then, CD11c+CD11b+ cells were gated following exclusion of doublets on a FSC-W/FSC-A plot. The gated cells were then further analyzed for antigen uptake or expression of CD86 or dectin-1. Representative density plots and histograms from an uptake experiment are shown. Grey histogram shows PBS control, dotted line indicates cells incubated with pHrodo-OVA and solid line cells incubated with pHrodo-LamOVA.

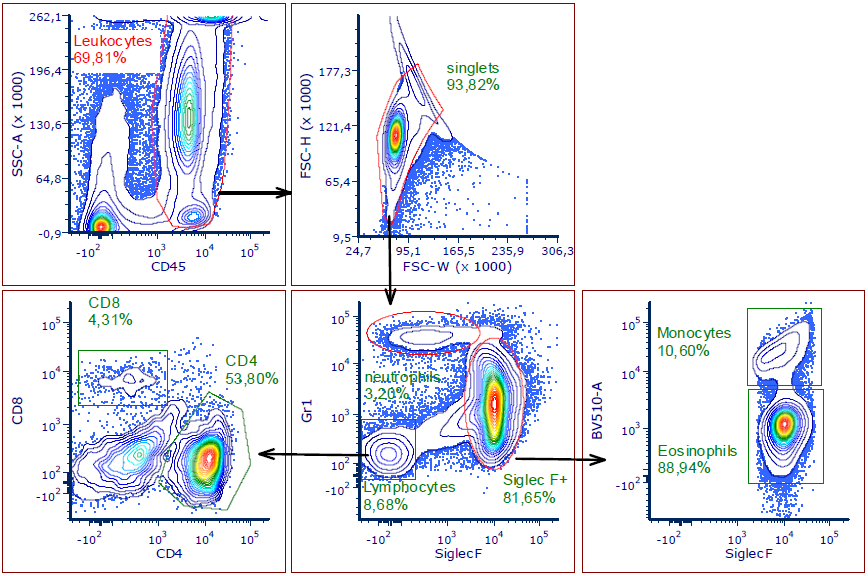

**Supplementary Figure 9. Gating strategy for BAL Leukocytes**. Total Leukocytes were gated based upon expression of CD45 followed by doublet exclusion. Neutrophils and lymphocytes were gated on a Siglec-F vs. Gr-1 plot. SiglecF+ cells were discriminated into monocytes and eosinophils based on autofluorescence on the BV510 channel. Lymphocytes were further distributed into CD4+ and CD8+ cells.

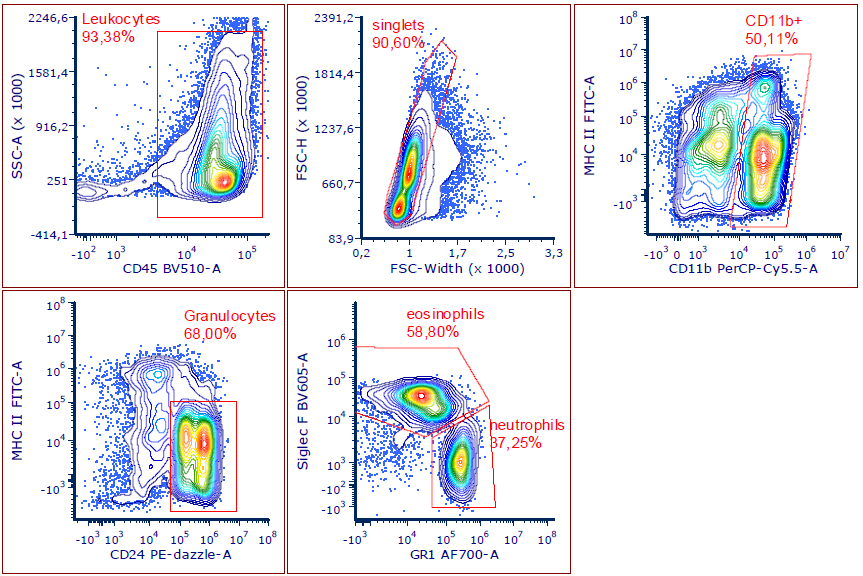

**Supplementary Figure 10. Gating strategy lung digests**. Total Leukocytes were gated based upon expression of CD45 followed by doublet exclusion. CD11b+ cells were gated on a MHC-II vs. CD11b plot. Granulocytes were identified as CD24+ MHC-II- cells and further separated into eosinophils and neutrophils based on SiglecF and Gr-1 expression.

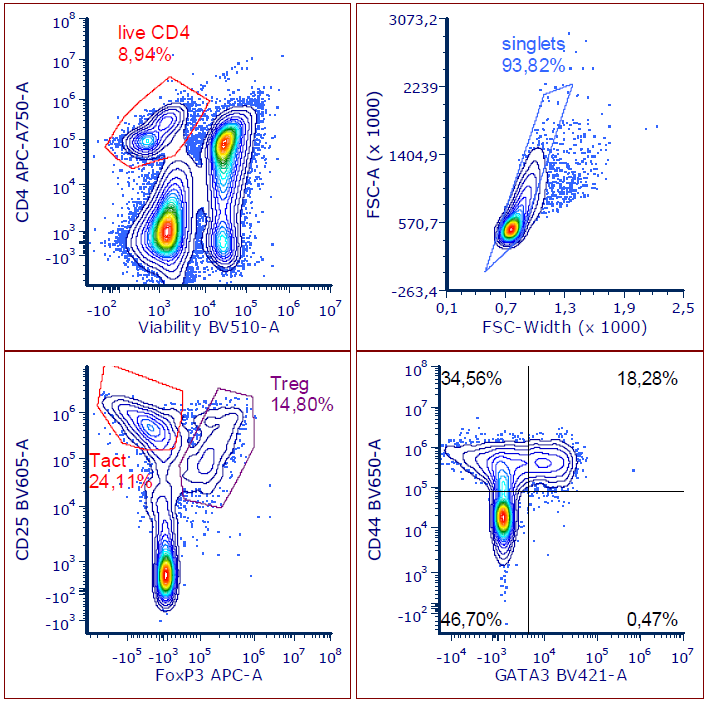

**Supplementary Figure 11. Gating strategy splenocyte cultures**. After in vitro restimulation, live CD4 cells were gated followed by doublet exclusion. Activated T cells (Tact) and regulatory T cells (Treg) were gated based on their expression of CD25 and FoxP3. Th2 cells were identified as CD44+GATA3+.
